# Small Data Challenge: Structural Analysis and Optimization of Convolutional Neural Networks with a Small Sample Size

**DOI:** 10.1101/402610

**Authors:** Rhett N. D’souza, Po-Yao Huang, Fang-Cheng Yeh

## Abstract

Deep neural networks have gained immense popularity in the Big Data problem; however, the availability of training samples can be relatively limited in certain application domains, particularly medical imaging, and consequently leading to overfitting problems. This “Small Data” challenge may need a mindset that is entirely different from the existing Big Data paradigm. Here, under the small data setting, we examined whether the network structure has a substantial influence on the performance and whether the optimal structure is predominantly determined by sample size or data nature. To this end, we listed all possible combinations of layers given an upper bound of the VC-dimension to study how structural hyperparameters affected the performance. Our results showed that structural optimization improved accuracy by 27.99%, 16.44%, and 13.11% over random selection for a sample size of 100, 500, and 1,000 in the MNIST dataset, respectively, suggesting that the importance of the network structure increases as the sample size becomes smaller. Furthermore, the optimal network structure was mostly determined by the data nature (photographic, calligraphic, or medical images), and less affected by the sample size, suggesting that the optimal network structure is data-driven, not sample size driven. After network structure optimization, the conventional convolutional neural network could achieve 91.13% in accuracy with only 500 samples, 93.66% in accuracy with only 1000 samples for the MNIST dataset and 94.10% in accuracy with only 3300 samples for the Mitosis (microscopic) dataset. These results indicate the primary importance of the network structure and the nature of the data in facing the Small Data challenge.

## Introduction

Deep neural networks have regained immense popularity in recent years. With the availability of much more powerful computational resources, such as CPU clusters and GPUs (Graphics Processing Units), studies are now capable of constructing network structures with even more and wider layers than before. The growth of deep neural networks from the 8 layer AlexNet[1], to the 19 layer VGG [2], to the 22 layer GoogleNet [3], followed by the 152 layer ResNet [4], shows a clear generalization of the idea that deeper networks perform better, and it has been pretty conclusive that the evolution trend of artificial neural networks is in the direction of even deeper or more complicated structures. However, this trend is ubiquitously built up-on the assumption of a sufficiently high sample size (i.e. Big Data), and the issue of overfitting or under-training can be reasonably ignored. While in numerous real-world applications, the number of samples in a dataset can be relatively limited, and the issues would arise with a generalized method of plain layer stacking in an attempt to improve performance. This “Small Data” challenge would call for a completely different mindset and approach from the existing “Big Data” one. Specifically, In the case of a “Small Data” challenge (e.g. sample < 5,000), the lines are still blurred as to whether the deeper we go, the better we perform or at what depth or width we achieve maximum accuracy. This small sample size issue is of particular interest when neural networks are applied to medical images, including MRI, CT, ultrasounds, and histopathology digital images, which often have limited sample size restricted by the availability of the patient’s population and experts’ labeling. Typical strategies used in training Big Data may not be readily applicable to these applications.

One possible solution to the small sample size problem is to use pre-trained networks [5] [6], also known as transfer learning. These approaches have gained popularity in many fields to handle the lack of significant samples in a dataset. The idea is to initialize the neural network with the weights trained in the related domain(s) and fine-tune the model with in-domain data. This approach provides a reasonable initial state and may speed up training when the two domains are close. There have been certain high performing solutions in the field of medical imaging with respect to the use of a pre-trained network [7] [8]. However in fields using medical images, where the input pixel format (e.g. Non-RGB data such as grayscale ultrasonic/MRI images or RGB+Depth) can be entirely different from conventional photographic images, where using pre-trained networks from other image domains implicitly hypothesize that the optimal network structure is *universal* (“Transferrable” networks) and not (or less) data-driven. Even if the pixel format is identical, whether a pre-trained network is a complete and reliable solution in the aforementioned fields needs to be rigorously examined, as in certain datasets in the cited articles, performance improvement using a pre-training set, if not substantial, could indicate a sizable similarity between the pre-training set and the target set, indicating that in many cases pre-training could only work if sets were substantially similar. There can also be a “Negative Transfer” in cases where there is a large domain gap between the pre-training set and the in-training dataset, resulting in a drop in performance, on the in-training after fine-tuning, compared to from-scratch training. Hence there still exists some ambiguity in the foolproof nature of the pre-trained network methodology.

Furthermore, as we encounter a need to apply neural networks to smaller and smaller sample size, the challenge is that we do not know whether a smaller subset of a larger set, may have the same optimal network as the whole dataset. For example, with a smaller size of training data, a very deep network (e.g. ResNet) may not generalize well since it may overfit the small training data requiring the need to apply a different learning scheme (e.g. adding regularization) or another strategy specialized for small datasets. Therefore, the questions are whether not only does the nature of the problem affect the optimal network, but also the size of the set, and how does this change in sample size affect the structure of the optimal network. This can be seen in fields (such as medical imaging) where the primary issue is the lack of a substantial amount of examples as well as extremely skewed classes in the training set and hence the ambiguity in the ad-hoc performance of the designed network. In addition, we do not know whether generalized network building rules, like “deeper the better” or “wider the better” will always work, specifically for smaller sized datasets. Since the rules are assuming that the dimension of the entire sample size is substantially greater than the network complexity.

Here we aim to address the above-mentioned questions when it comes to training a convolutional neural network with a small sample size. First, we exhaustively iterated a list of networks given a upper limit of the complexity, trained them, and studied their performance, aiming to understand the dependence of the performance of the network for each dataset on the structural hyperparameters, namely the nature of layers (e.g. convolution, fully connected, max-pooling…etc.), number of layers and the dimension of layers. Next, we examined the “nature versus size” hypothesis: whether the optimal structure of the neural network, under the identical pixel format (RGB), is largely determined by the “nature” of the data (e.g. photographic, calligraphic, microscopic images) or the “size” of the data. In other words, we would like to study whether datasets from different data sources or image modalities would have very similar or different from the optimally performing networks, in comparison with the influence of a different sample size on the optimal network structure. If the effect of “data size” is greater than the effect of “data nature”, then it means the optimal network will converge to a complicated network structure as data size become larger, and we can ignore the issue that a large complicated pre-trained network is originally from a different image modality and always use a more complicated network if the pixel format is identical. On the contrary, if the effect of data nature is greater, then the use of a pre-trained network from different image sources may not be favored. Moreover, it will support a strategy that, when it comes to a Big Data problem, we can logically use only a subset of it to optimize the network structure. Through the study of this hypothesis we aim to validate or invalidate the ability of pre-trained network as a universal watertight procedure in the field of smaller sample sizes (medical imaging, etc), against the opposite approach that will require a search for the best performing network specifically designed for smaller-sized datasets, which we have established as an issue in various fields. Lastly, we will calculate the best performance a convolutional neural network can achieve after optimizing the network structure and compare it with the average performance from randomly selected networks.

To examine this, we used three image datasets from entirely different data nature, including MNIST(calligraphic) [9], CIFAR10 (photographic) [10], and Mitosis image data (microscopic). We trained and tested each structure followed by layer dimension (layer width) optimization using small subsets (less than 5,000 samples) of these datasets, and investigated the performance difference in different network structures, as detailed in the following sections.

## Materials and Methods

Given a limited size of training samples, we searched a constrained space of networks in two iteration rounds. First, we generated a list which contains all the possible network structures with a fixed dimensionality of layers (e.g. channel size of a convolutional layer), given a particular constraint which we set based on using the network’s Vapnik–Chervonenki (VC)-dimension [11] [12] [13]. We then set a maximum value of the VC-dimension and hence generated a list of structures with VC-dimension less than or equal to the maximum value. The list of structures was generated recursively using a tree data structure, where the paths of the tree from the root to the leaves each denote a network structure in this constrained space. All networks were trained and tested with a held-out validation set to record their accuracy, thereby allowing us to study the relation of network accuracy to different network structures, as well as select the best-performing ones for each step. Second, we selected 5 optimally performing networks from the first round to generate similar networks with different layer dimensions. Specifically, we iterated over possible permutations of layer dimensions to find the optimal set of dimensions for the optimal structure found in the first step. All networks were also trained and tested on the held-out validation set to study their performance. The exact way in which we defined and calculated the VC-dimension, how we generated the tree structure and how the network structures/dimension permutations were trained, validated and tested is elaborated as follows:

### Generating Network Structures and Layer Dimension Permutations

Based on the input dimension of the images in the dataset and the number of structures deemed suitable for our analysis and the testing of our hypothesis, we set a maximum value for constraining the complexity of networks which can be generated. This value reflects the “learning capacity” of each network structure. Specifically, we calculated the maximum points of data the network could shatter (divide) using the VC dimension of the network as a measure of the complexity of the network (Vapnik, et al. 1994) (Sontag 1998).

The nearly-tight bound of the VC dimension for any multilayer neural network, with a piecewise linear function was used as calculated in the paper by Harvey, Liaw and Mehrabian (Harvey, et al. 2017). We used the absolute upper bound to limit the size of a network. It is expressed as:

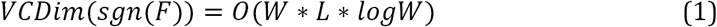

Where *W* is the total number of weights in a network (including bias), and *L* is the number of layers in the network in which these weights are arranged. *F* is the set of functions (real-valued) computed by this network. The dimension of the image at any layer is (*n* × *n*) where *n* is the number of pixels in the respective dimension. All windowing/filter based operations (pooling and convolution) had the same padding that is, adding zeros to the edges to match the size of the filter (*k*) (this technique may be extended to other padding techniques as well).

For any convolutional layer, the total number of weights were computed as:

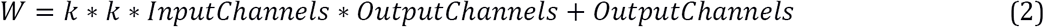

For any fully-connected (dense) layer the total number of weights are computed as:

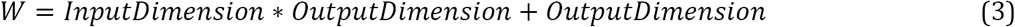

Max-pooling layers do not have any weights, and hence do not contribute to the VC-dimension (they, however, change the dimension of the input for the following layers and hence do play a role in affecting the VC-dimension of the entire network structure). We sum up all the values of the weights of all the layers and then apply the formula for the VC-dimension. For the purpose of building the recursive tree structure and terminating a branch, we keep an upper bound on the VC-dimension of any network, called the ‘maximum VC-dimension’. This value can be greater or smaller based on what is required for the procedure’s application.

To construct a list of eligible network structures, we made use of a tree data structure to topologically arrange the various possible layers of a fixed dimension/width. At each node of the tree, we calculated the VC-dimension (with the corresponding final output layer added) of the structure. We checked if the VC-dimension is lesser than or equal to the maximum allowed VC-dimension. We then placed the node and recursively called the building function. This node could contain either a Convolutional Layer, Maximum Pooling Layer, a Fully-Connected (Dense) Layer or the final Output Layer. At the root of the tree, we necessarily had a fixed input dimension layer. The first layer could be any of the above possible layers, however, the input dimensionality of a Fully Connected layer would be that of the input sample and the number of input channels of the Convolutional Layer could be strictly equal to the number of channels in the input image.

If the maximum VC-dimension is not greater than the value of the VC-dimension of the network with the last fully connected layer, we constructed the last layer and placed a terminating leaf at that point or we added in a new layer and continued that branch of the tree. After we add in a layer, we recursively keep track of the total number of weights and layers in which the weights are arranged, to keep track of the VC dimension of the possible network structures at each layer. In that way we recursively generated a tree structure which contains nodes from which emerging branches represented all the possibilities of the layers, which satisfied the maximum VC-dimension value condition such that when we enlist all possible paths from the root of the tree to all the leaves, it represented all the possible network structures (which could be created with all the possible layers) which have a total VC-dimension value lesser or equal to the overall maximum VC-dimension constraint. We limited the possible filter size of the convolutional layers to 5×5 or 7×7, the max-pooling layers to have a pooling-window size of 2×2 or 4×4 and a fixed layer channel dimension of 10.

We used a maximum VC-dimension of 3,500,000 for MNIST and CIFAR10 (taking into consideration the RGB channels). A total of 7103 different network structures were generated for the MNIST input dimension and 2821 different network structures for the CIFAR10 input dimension. The mitosis dataset has a different input image dimension, and we used a maximum VC-dimension of 2,725,000 and created a list of 2599 network structure proposals. We trained and tested all networks structures and recorded each of their performance to examine whether the optimally performing structure is largely affected by the data nature or data size.

The second iteration studies the performance in different layer dimension (or width). We first pick the 5 best performing ones (lowest error) and permuted all possible combinations of layer dimension to generate a second list of networks. These networks shared the same structural configuration, but with different possible combinations of their layer dimensions. We allowed the possible dimensions of each layer to grow in powers of 2, i.e. 32, 64, 128, and so on. Since the depth and nature of each optimal set of structures are different, the number of generated layer dimension permutations differs for different dataset/subset combinations.

All code for network structure list generation, training, validation, and testing can be found at: (https://github.com/rhettdsouza13/DNN-And-HDFT.git)

### Training/Validation Method and Datasets - MNIST

The MNIST [9] handwritten digit recognition image dataset contains 60,000 training image samples and 10,000 test image samples. We randomly selected 1000, 500 and 100 samples from the 60,000 samples as the possible training sets. We then made the following division for training/validation. The 1000-sized set was divided into 800 samples for training and 200 samples for validation. The 500-sized set was divided into 400 samples for training and 100 samples for validation. The 100-sized set was divided into 60 samples for training and 40 samples for validation.

We made use of the ADAM method of Stochastic Optimization [14] with 100-sized mini-batches for the cases of the 1000 and 500 data samples and 10-sized mini-batches for the case of the 100 data samples. We made use of a softmax cross-entropy with logits objective function. The learning rate was kept constant at the recommended value of 0.001 for all epochs of training. No weight decay was used. A total of 8, 4 and 6 iterations of the gradient descent of the ADAM optimizer per epoch, were used for 1000, 500, and 100 datasets respectively. After every epoch of training, we validated the model.

We implemented an early stopping protocol to avoid overfitting and reduce the time of training [15]. After each epoch, we checked the validation error and calculated if it was the minimum. If the network didn’t reduce its error in the next 5 epochs, we stopped training. This signified either one of two things. Either the network had reached an optimal minimal error value, or the error was beginning to increase and it was overfitting. We recorded the value of the validation accuracy and validation error, for each epoch.

To generate all possible combination of network structures, we employed the tree building technique to exhaustively list all the possible layer combinations. We allowed different filters of the convolutional and max-pooling layers to represent different structures, and for each network structure, we assigned four possible constant values (10, 20, 40, 80) to the channels of the convolutional layers and also the output dimension of fully connected layers. We generated a total of 7103 different network structures. We then trained the networks using the technique mentioned in the previous section and ranked them according to their classification error in their validation curves, representing the fully-trained state of the network/structure and selected the networks with the 5 lowest errors to be used in the next step. For these 5 best-performing networks, we further permuted their channel dimensionality, to find the optimal configuration of network channel dimensions. Keeping everything else constant, the value of the network dimensions was allowed 3 possibilities for demonstration purposes, 32, 64 and 128. We generated 9261 networks for the 100-sized subset, 3267 networks for the 500-sized subset and 13729 networks for the 1000-sized subset. We then trained the various dimension configurations using the techniques mentioned in the training sections and then ranked them in a similar fashion as the previous step.

### Training/Validation Method and Datasets - CIFAR-10

We applied the same optimization approaches to CIFAR-10. The CIFAR-10 [10] dataset consists of 60,000, 32×32 color images (RGB) in 10 classes, with 6,000 images per class. The original dataset has a total of 50,000 training images and 10,000 test images. Here we only used 5,000 samples. In details, we extracted 5000 random samples from the 50000 training set. This 5000 was further subdivided into 4000 for training, while 1000 examples were kept aside for validation for each epoch. The optimizer settings used in the MNIST dataset were used here as well. In the case of CIFAR-10, a total of 40 iterations of gradient descent of the ADAM optimizer per epoch were run, for the 4000 sized training set. After every epoch of training, we validate the model with the 1000-sized validation set. We applied the same early stopping mechanism as earlier mentioned. We recorded the value of the validation accuracy and validation error, after every epoch.

Then, like MNIST, we iterated overall network structures and trained them using the techniques mentioned in the training sections and then ranked them in a similar fashion as the previous step. After ranking the networks like with the MNIST dataset, we generated their layer dimension permutations with dimension possibilities of 32, 64, 128 and 256. We generated a total of 1172 different layer dimension combinations for the top 5 network structures from the previous step.

### Training/Validation Method and Datasets - Mitosis

The mitosis dataset is from the Tumor Proliferation Assessment Challenge 2016 (TUPAC16, MICCAI Grand Challenge, http://tupac.tue-image.nl/node/3). The data consists of images from 73 breast cancer cases from three pathology centers, with 23 cases previously released as part of the AMIDA13 challenge. These cases were originally from the Department of Pathology at the University Medical Center in Utrecht, The Netherlands. The remaining 50 cases were collected from two different pathology centers in The Netherlands. The slides were stained by H&E stains, and whole-slide images were produced with the Leica SCN400 whole-slide image scanner at ×40 magnification, resulting in a spatial resolution of 0.25 μm/pixel. Each case is represented by a number of image regions stored as TIFF images.

The location of a positive finding of mitotic cells was annotated by at least two pathologists. The negative labels were generated using WS-recognizer (http://ws-recognizer.labsolver.org). The tool first extracted stains color from mitotic cells to recognize other targets with similar stains in the histology images [16]. The negative targets were thus generated by excluding the pathologists’ labels from targets recognized by WS-recognizer. Both positive and negative sample were cropped from the original TIFF images as a 64×64 RGB image. The resulting dataset contains a total of 4290 samples. Examples of the microscopic images can be seen in Fig. 1.

**Fig. 1.**
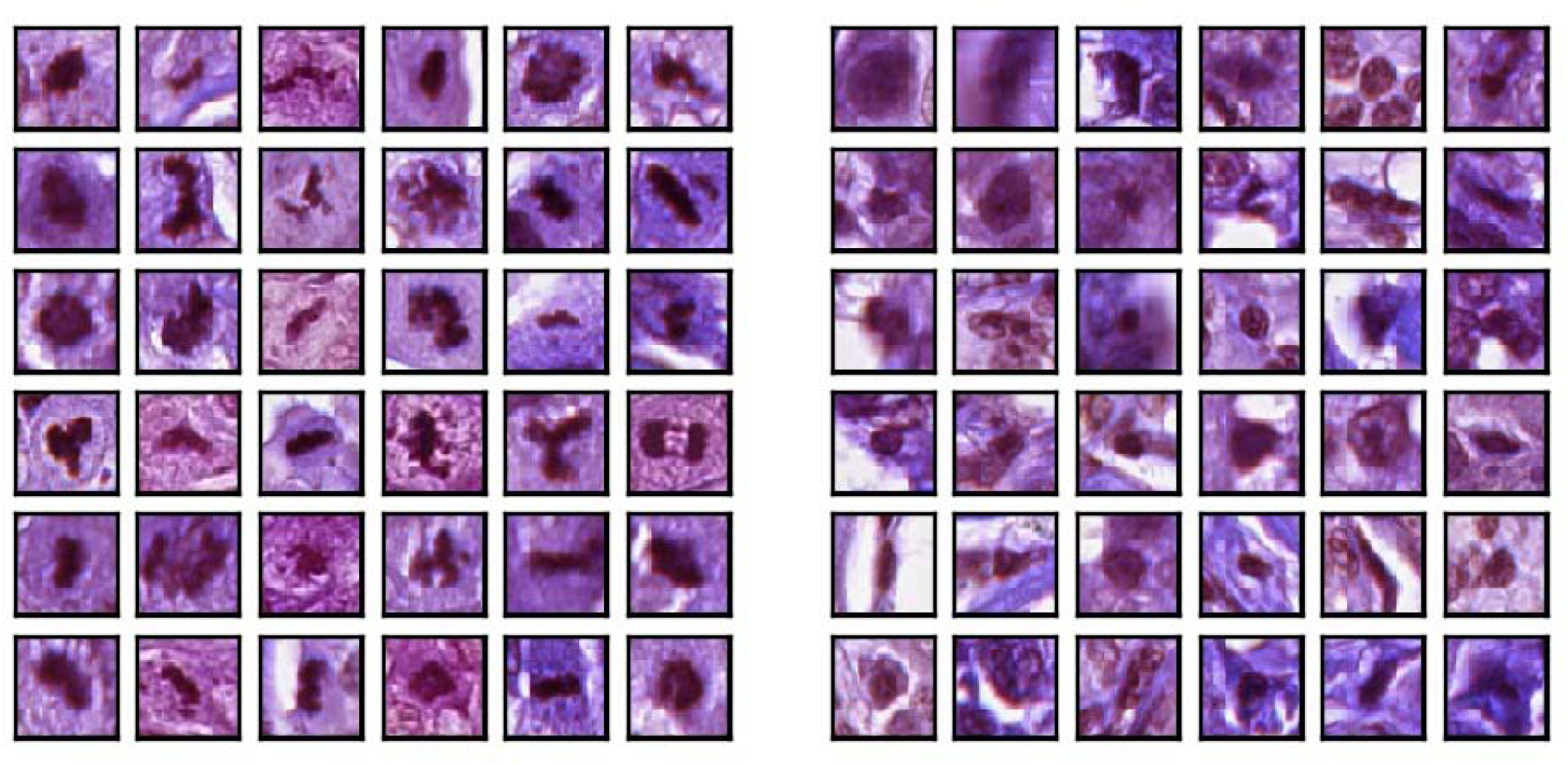
The mitosis dataset used in this study showing positive samples (left), and negative samples (right). The negative samples were extracted using an automatic stain recognition tool. The overall dataset contains a total of 1430 positive and 2860 negative samples.

To test our hypothesis, the dataset was divided into three sets for the sake of training, validation, and testing of sizes 2500, 800 and 990 respectively. This set represents a real-life situation in a field like medical imaging where the number of examples is limited. The optimizer settings used in the MNIST/CIFAR-10 dataset was used here as well. Therefore, In the case of the Mitosis dataset, a total of 25 iterations ran for the gradient descent section of the ADAM optimizer per epoch, for the 2500 sized training set. After every epoch of training, we validate the model with the 800-sized validation set (initially kept aside from the 3300). We applied the same early stopping mechanism as earlier mentioned. We recorded the value of the validation accuracy and validation error, after every epoch.Then, as performed for the earlier datasets, for the sake of the other dimension cases, we replace the 10 dimension with, 20, 40 and 80, to get the same structures but with channel dimensionality of the corresponding values respectively, to remove any bias towards the fixed layer dimension. We then trained the structures using the technique mentioned in the previous section and ranked them according to their classification error in their validation curves, representing the fully trained state of the network/structure and selected the networks with the 5 lowest errors to be used in the next step.

Like earlier with MNIST/CIFAR-10 dataset, once we found the set of optimal structures, we then permuted their channel dimensionality, to find the optimal configuration of network channel dimensions. Keeping everything else constant, the value of the network dimensions were allowed 2 possibilities for demonstration purposes, 32 and 64, and consequently generated 576 different possible networks. More and/or different values may be used, based on the requirement and application. We then trained the networks using the techniques mentioned in the training sections.

## Results

### Frequency Distribution of Classification Error

Fig. 2 and Fig. 3 are the histograms displaying the distribution of the validation error rate for the different dataset-sample size, namely CIFAR-10 (5000 samples), MNIST (1000 sample) and Mitosis (3300 sample) (Fig. 2) and MNIST (100 sample), MNIST (500 sample) and MNIST (1000 sample) (Fig. 3). The X-axis represents the classification error (in %) of the network structures and the Y-axis represents the number of networks in the said error rate range. The classification error on the X-axis is divided into a bin size of 2%,

**Fig. 2.**
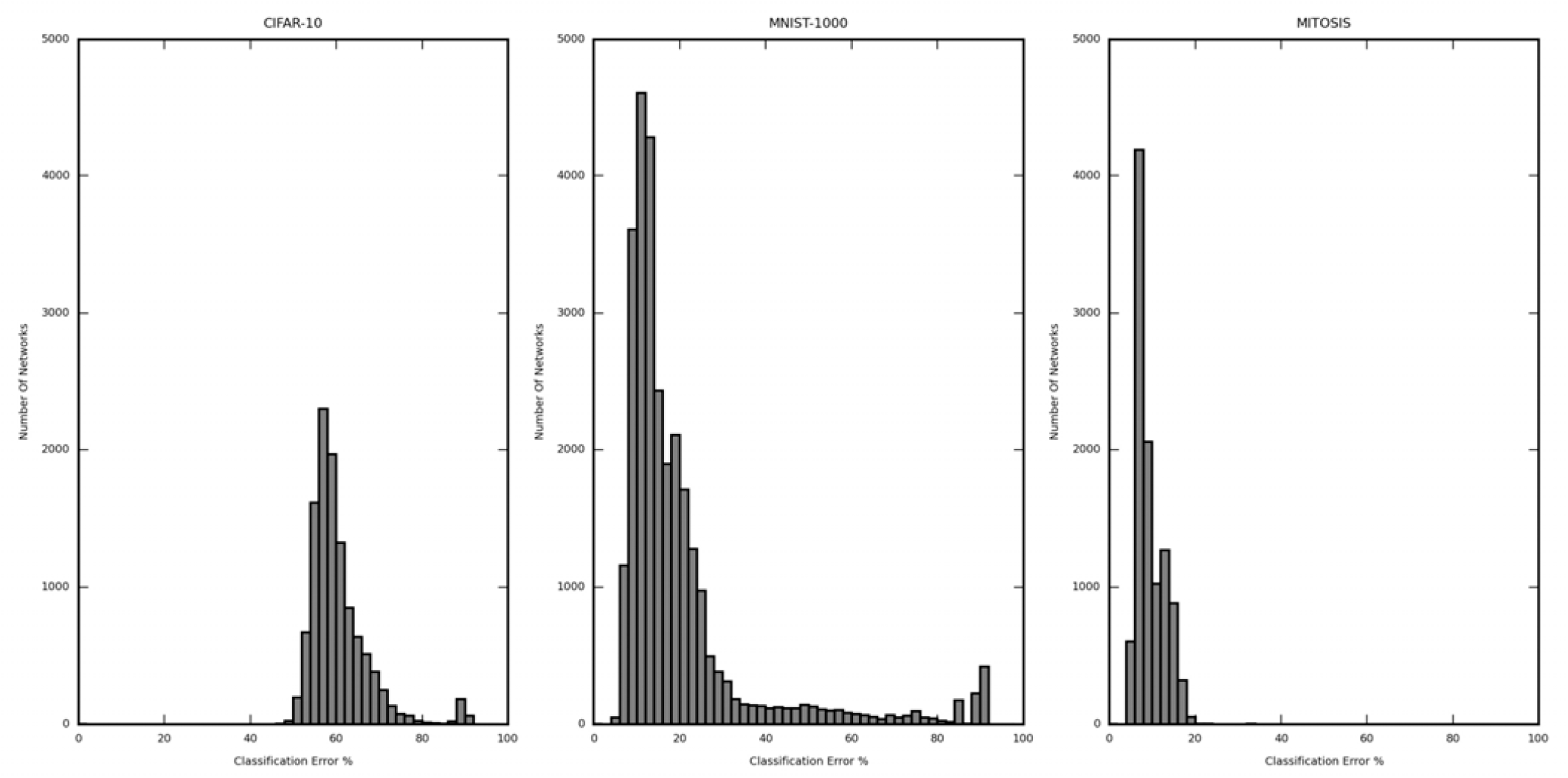
Histogram showing the distribution of the classification error from all iterated network structures trained by 1000-sized CIFAR10, 1000-size MNIST, and Mitosis dataset. The distribution differs substantially in different datasets and for the same dataset, different structures may yield very different error rates.

**Fig. 3.**
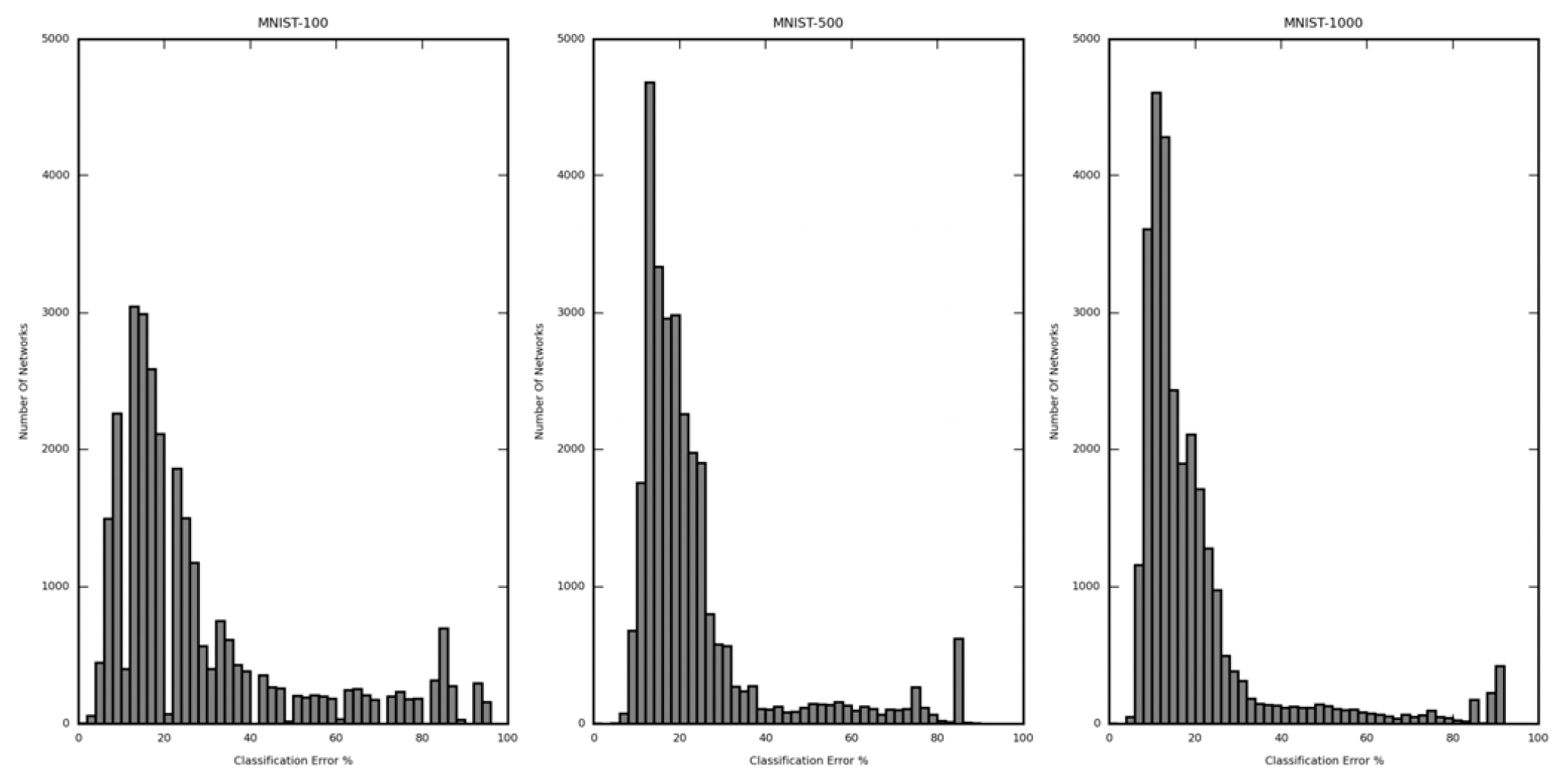
Histogram showing the distribution of the classification error from all iterated network structures in each subset of the MNIST dataset (100, 500 and 1000 samples). The three distributions are similar regardless of sample size, suggesting that the sample size contributes less to the optimal network structure search procedure.

Fig. 2 and Fig. 3 show the frequency distribution of the lowest per network validation classification error. The difference in performance between the best and the worst model is considerable (in later comparison to Fig. 6 and Fig. 7). In Fig. 2, the calculated (per network lowest) average classification error for CIFAR-10 (5000 samples) is 60.49%, MNIST (1000 samples) is 20.11% and Mitosis (3300 samples) is 9.43%. The classification error for the best performing networks is 47.80%, 7.00%, and 7.37% respectively. The improvement of the best performing network structures over the average case was 12.69%, 13.11%, and 2.06% respectively.

In Fig. 3, the similar analysis for different subsets of the MNIST dataset, also shows that the difference between the largest and the lowest classification error is considerable. This means that the number of values of error taken by the networks is large enough to justify the selection of the best network. The calculated average classification error for MNIST (100 samples) is 27.99%, MNIST (500 samples) is 23.44% and MNIST (1000 samples) is 20.11%. The classification error is 0.00% (validated against only 40 samples, further explanation in subsequent sections), 7.00% and 7.00% respectively. The improvement of the best performing network structures over the average case was 27.99%, 16.44%, and 13.11% respectively.

### Effect of Network Structure on Classification Error

In Fig. 4 the lowest validation classification error (in percentage) for each structures’ validation curve has been plotted against various important network characteristics and attributes, for each dataset-sample size, namely CIFAR-10 (5000 sample), MNIST (1000 sample) and Mitosis (3300 sample).

**Fig. 4.**
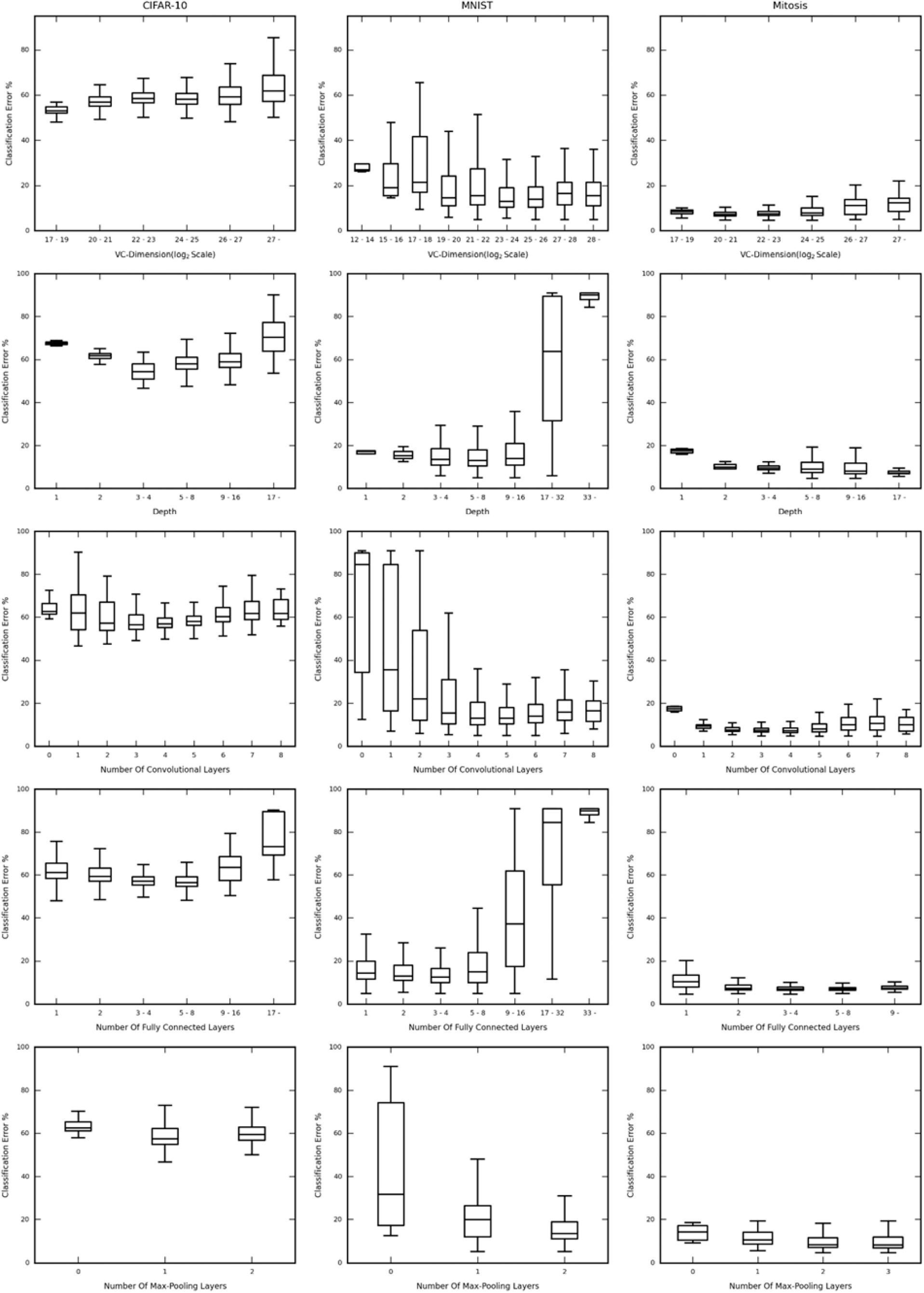
Box and whisker plots showing the distribution of the classification error against the structural hyperparameters (i.e. VC-dimension, Depth, Number of Convolutional Layers, Number of Fully Connected Layers and Number of Max-Pooling Layers) for subsampled CIFAR-10, MNIST and the Mitosis dataset (columns). Notice the difference in the trends followed for each dataset, indicating that the effect of the structural hyperparameters on the performance of each structure, for each dataset is unique. Also note the difference in the values of the structural attributes at which the overall lowest validation classification error is achieved, indicating a unique optimal structure for each dataset.

The figure consists of 3 columns, for each dataset (mentioned above). Each row contains the classification error percentage box and whisker plot v/s the VC-dimension (log scale), depth, number of Convolutional Layers, Number of Fully Connected Layers and Number of Max-Pooling Layers respectively. In cases where the range of values are too large, we have binned those values for better graphical representation and clarity. The networks have been binned based on their VC-Dimension, Depth, and Number of Fully Connected Layers, in the associated box and whisker plots. However, Number of Convolutional Layers and Number Of Max-Pooling layers have been shown individually due to the lower number of possible values taken by those attributes. The ranges of the Y-axes’ (Classification Error %) have been kept constant across each row. This is to provide efficient comparative analysis, across each dataset.

In general, the best performing networks of each dataset, are very different from each other, in other words, the most optimal network structure is data-driven. This means that a standard layer-to-layer building procedure for a network if used for all datasets, will very probably lead to a sub-optimal network being selected, as both the order and the number of the corresponding layers play a part in deciding the performance of the network for each specific dataset.

The result also suggests that “deeper the better” is not always true for conventional convolutional neural networks for small datasets. In most cases, the performance degrades quite drastically as the depth increases. The plots clearly show, that as we increase the depth there is an initial drop in the classification error, but the error soon rises sharply (CIFAR-10 and MNIST). However, this may not always be true, as we can see that the Mitosis dataset has no clear bias toward deeper or shallower networks.

When discussing the individual layer characteristics, as in, Convolutional, Fully-Connected or Max-Pooling Layers, we can see that each dataset has a specific optimal number of layers for each category. Empirically, our architectural search may iteratively find the feasible classification network for the given dataset. However, the reason why there is no generalizable network design strategy is unknown and is likely to be data-driven. We left the theoretical approach to disentangle such myth as our future work.

In Fig. 5 the classification error (in percentage) is plotted for different sample sizes of the MNIST dataset, namely 100, 500 and 1000 sized sets against the various network attributes, in the same way as in Fig. 4. The result shows high consistency in these plots (across a row). This is particularly interesting as this may suggest that the optimal network structures are driven primarily by the nature of the data. The smaller differences displayed from sample-size to sample-size can still be viewed, leading to minute differences in the optimal structure for each sample. One small preference can be seen is that, the larger the set, the deeper is the preference, until an optimal point, after which performance degrades. However, the nature of the layer added in, also plays an important role, as in; there is a common positive correlation between a number of fully connected layers and classification error, hinting that “deeper the better” may not hold true if the layers being stacked on are fully-connected layers.

**Fig. 5.**
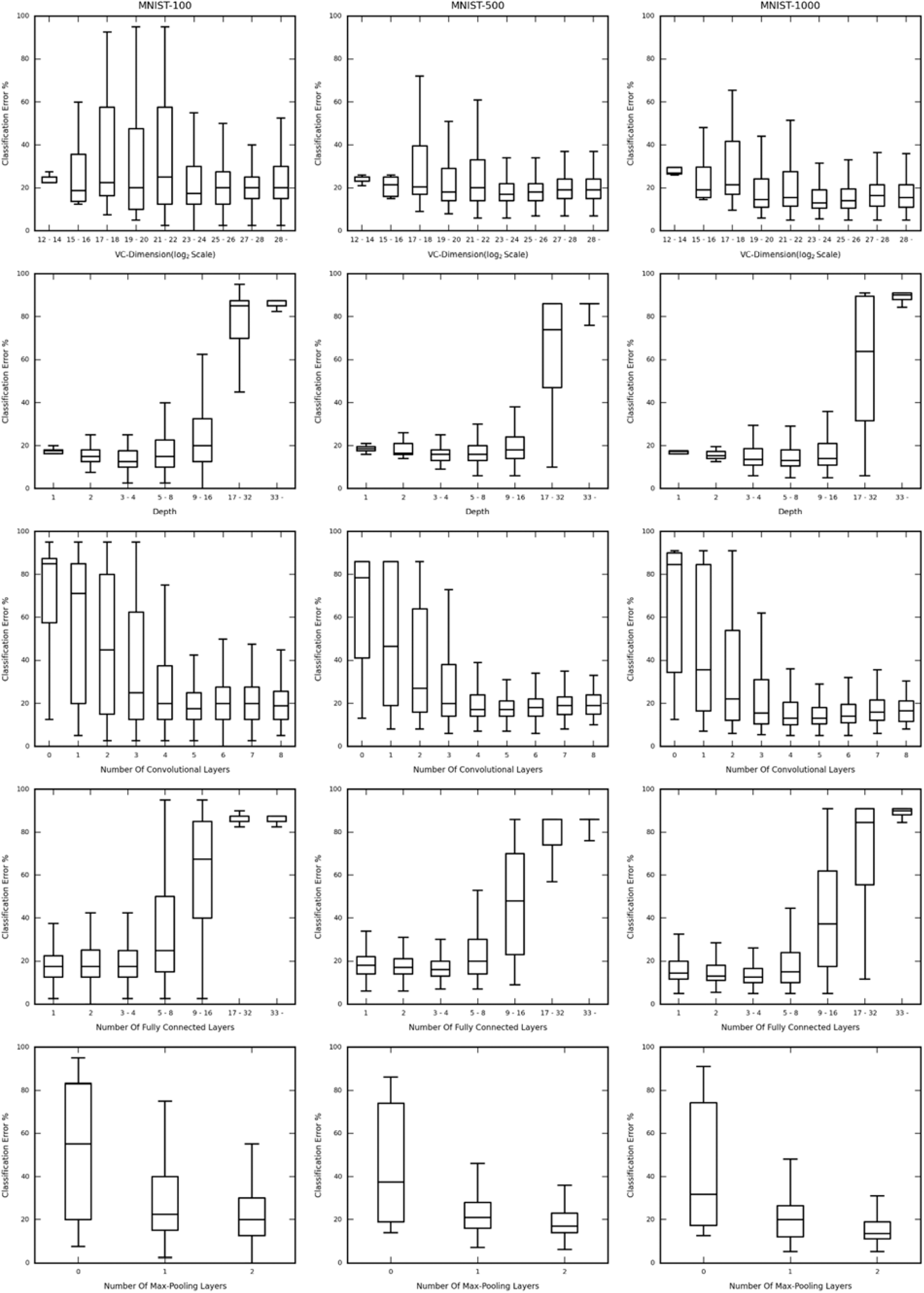
Box and whisker plots showing the classification error for each network structure against the structural hyperparameters (i.e. VC-dimension, Depth, Number of Convolutional Layers, Number of Fully Connected Layers and Number of Max-Pooling Layers) for MNIST dataset of different sample size. Notice the similarity in the trends followed for each subset, indicating that the effect of the structural attributes on the performance of each structure, for each subset is similar, indicating that, the optimally performing structures is highly data-driven.

Another implication is that if the trends in the same dataset are common, we may operate only on a smaller subset of the larger dataset and obtain a reasonably high performing structure for the larger dataset as well. This can greatly reduce the time required to train and validate the list of the various different structures or attempts to manually construct a high performing network. As a disclaimer, fine-tuning of the structure may be required for the larger set as sample size may play a role in affecting the optimal number of layers. Nevertheless, the main process will generate a reasonably good starting point.

### Optimal Network Structure

Table 1 lists the best performing structure obtained from the training and validation procedure mentioned earlier for each dataset/sample-size combination. The highest cross-validation accuracy calculated was 92.63% for the Mitosis dataset, 52.20% for the CIFAR-10 dataset, 93.00% for the MNIST: 1000-sized set, 93.00% for the MNIST: 500-sized set and 100.00% for the MNIST: 100-sized set, whereas the classification accuracy that was calculated from the full test set of the original dataset, was 92.93%, 50.46%, 92.68%, 85.34% and 60.16% respectively.

**Table 1.**
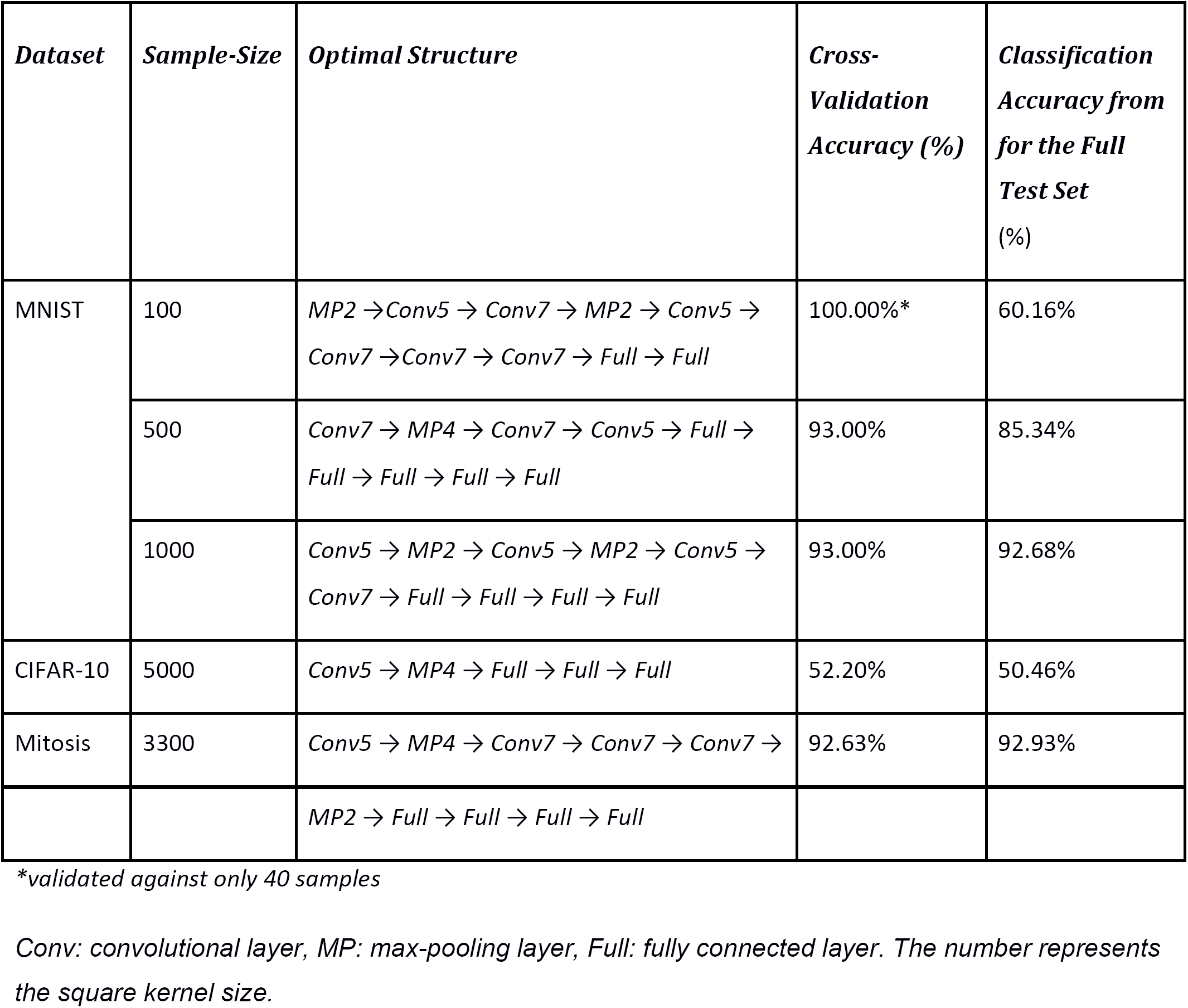
This table presents the results of the structure selection step.

A major observation we can make is in agreement with the earlier section from Fig. 4 and Fig. 5, is that the best performing structures (without taking channel width/dimension into consideration) are widely different for different datasets. One perceivable issue is the presence of the 100% validation accuracy for the MNIST-100 sample. This is because, when dealing with an extremely small sample like 100 with a split 60-40 for training-validation sets, we can have a substantial probability of the network easily getting all of the 40 examples correct or wrong. This can be avoided using different cross-validation techniques. However, to keep the comparison valid between all the various sets and subsets, we had to keep the techniques and training parameters uniform, to avoid any other factor influencing our choice of the optimal structure and attributing differing results to any arbitrarily distinguished procedure.

### Effect of Layer Dimension on Classification Error

Fig. 6 and Fig. 7 we use the histograms for the layer dimension permutation selection step, as we did for the previous step (structure selection). The axes have the same representation and meaning as earlier. The classification errors have the same meaning as earlier.

**Fig. 6.**
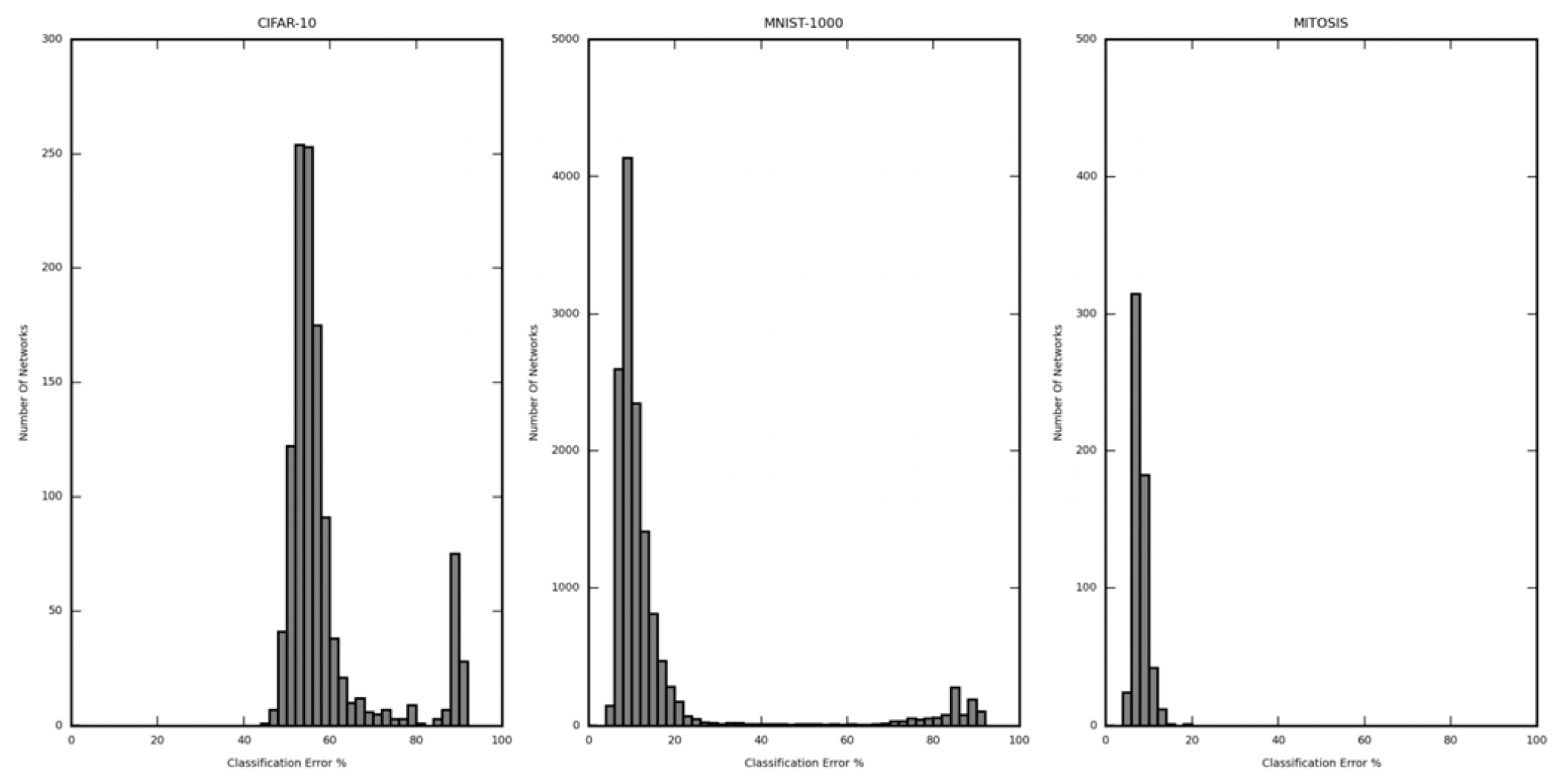
Histogram showing the distribution of classification error in the layer dimension optimization step for each dataset (subsampled CIFAR-10, MNIST, and Mitosis) The performance variation due to layer dimension is smaller than the structural optimization step and does not have a large difference among different width parameter. This suggests that the structural hyperparameters of the network, play a comparatively more dramatic effect on the performance of the network, than the layer width.

**Fig. 7.**
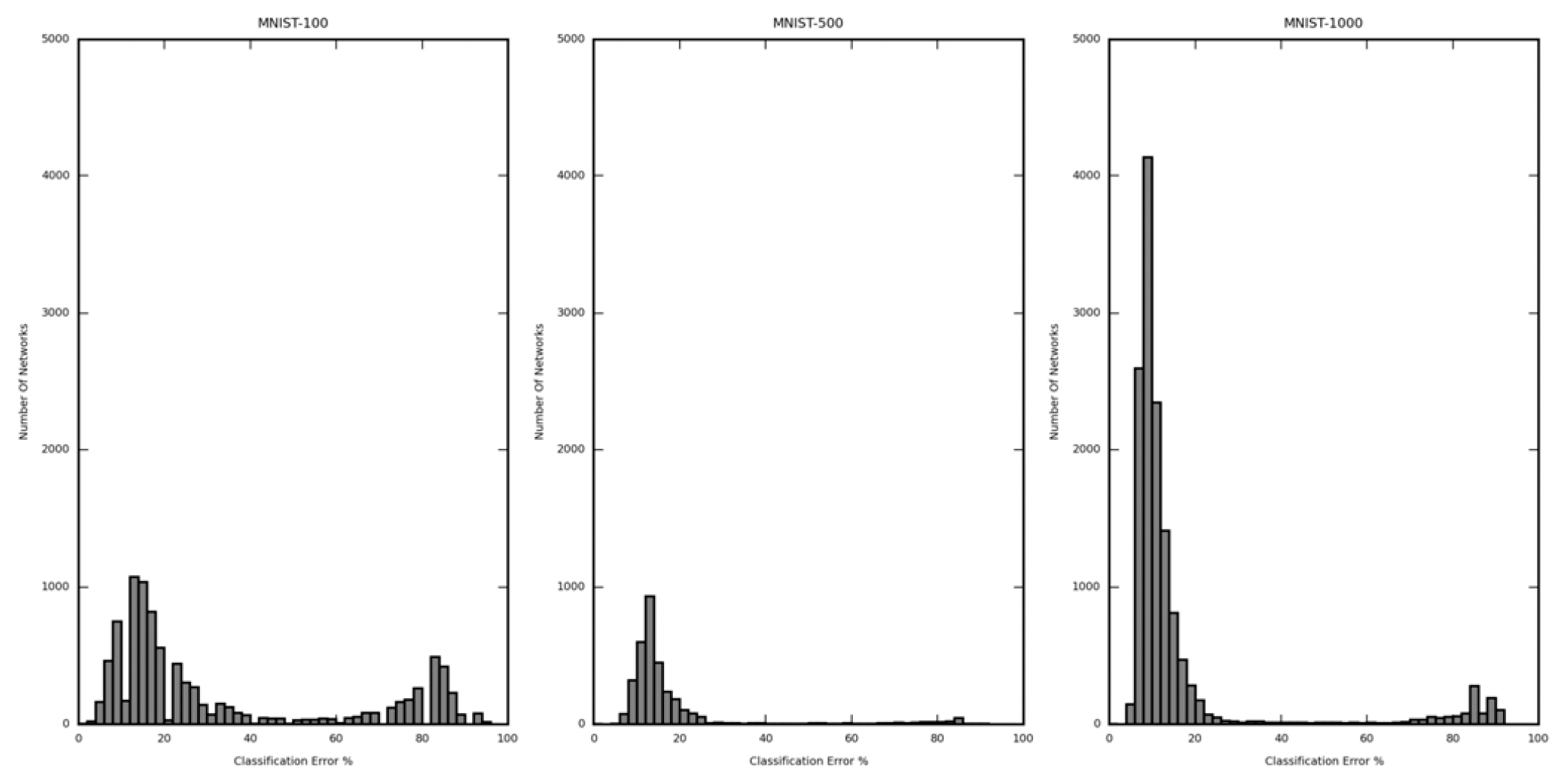
Histogram showing the distribution of classification error in the layer dimension optimization step for each subset of the MNIST dataset (100, 500 and 1000 samples). The performance variation does not differ a lot due to sample size.

A conclusion we can make is clearly the range which the networks occupy is much more narrow (not negligible) compared to the structure selection step. The calculated (per network lowest) average classification error for CIFAR-10 (5000 samples) is 58.88%, MNIST (1000 samples) is 16.58% and Mitosis (3300 samples) is 7.93%. The classification error for the best performing networks is 46.50%, 5.50%, and 4.13% respectively. The improvement of the best performing layer dimension permuted networks over the average case was 12.38%, 11.08%, and 3.8% respectively. This shows that the CIFAR-10 dataset and MNIST have a relatively higher dependence on the structure of a set, rather than the configuration of the layer dimensions. The Mitosis dataset seems to improve more dramatically over the average for the layer dimension optimization step than the previous step. However, the narrow range occupied by the histograms in Fig. 6 for the Mitosis dataset indicate that the performance of the networks is more influenced by the structure (wider range as observed in Fig. 2), than the layer dimension configuration. The same can also be viewed in the histograms of the CIFAR-10 and MNIST dataset subsets in Fig. 6 (histogram range).

The analysis for different subsets of the MNIST dataset, reveals the same conclusion. The calculated average classification error for MNIST (100 samples) is 33.70%, MNIST (500 samples) is 17.71% and MNIST (1000 samples) is 16.58%. The classification error is 0.00% (*validated against only 40 samples, further explanation in subsequent sections), 7.00% and 5.50% respectively. The improvement of the best performing layer dimension permuted networks over the average case was 33.66%, 10.71%, and 11.08% respectively. We can see in the case of the MNIST, 100 sample size case the average classification error rises for the layer dimension permutation step and hence the difference between the best and average case also increases. This can indicate a substantial dependence of the performance of the network on the width/channel dimension of its individual layers. On further analysis of the histogram in Fig. 7, qualitatively, we can say that for the MNIST, 100 sample size set, the networks’ performances are very sensitive to the nature of the specific permutation (channel dimension). However, since the final accuracy on the official MNIST 10,000-sized test set showed a rise in accuracy (the difference between full test accuracy in Table 1. and Table 2. for the MNIST, 100 sized subsets), we may conclude that this step did indeed result in an increase in the performance of the network.

**Table 2.** This table presents the results of the Layer Dimension Permutation step.

| ***Dataset*** | ***Sample-Size*** | ***Optimal Permutation*** | ***Cross-Validation Accuracy (%)*** | ***Classification Accuracy from for the Full Test Set (%)*** |
| --- | --- | --- | --- | --- |
| MNIST | 100 | *Conv5,64 → MP4,64 → Conv5,128 → Conv5,64 → Conv5,64 → Conv7,32 → Conv5,128 → Full,32 → Full,32 → Full,10* | 100.00%* | 68.33% |
|  | 500 | *Conv7,64 → MP4,64 → Conv7,128 → Conv5,64 → Full,128 → Full,128 → Full,128 → Full,32 → Full,10* | 93.00% | 91.33% |
|  | 1000 | *Conv5,32 → MP2,32 → Conv5,32 → MP2,32 → Conv5,128 → Conv7,32 → Full,64 → Full,64 → Full,32 → Full,10* | 94.50% | 93.66% |
| CIFAR-10 | 5000 | *Conv5,64 → MP4,64 → Full,64 → Full,32 → Full,10* | 53.50% | 55.18% |
| Mitosis | 3300 | *Conv5,32 → MP2,32 → Conv7,32 → MP2,32 → Conv5,64 → Conv5,32 → MP2,32 → Conv7,32 → Full,64 → Full,32 → Full,2* | 95.87% | 94.10% |

*\*validated against only 40 samples*
*conv: convolutional layer, mp: max-pooling layer, full: fully connected layer. The number represents the kernel size and layer width.*

### Optimal Layer Dimension Permutation

In Table 2, we can see the highest performing networks and their associated highest validation accuracies after the layer dimension permutation step obtained from the training and validation procedure mentioned earlier for each dataset/sample-size combination. The shorthand notation is the same as that in Table 1, with the only difference being the dimensionality of each layer has been written alongside. The highest cross-validation accuracy calculated was 95.87% for the Mitosis dataset, 53.50% for the CIFAR-10 dataset, 94.50% for the MNIST: 1000-sized set, 93.00% for the MNIST: 500-sized set and 100.00% for the MNIST: 100-sized set, whereas the classification accuracy that was calculated from the full test set of the original dataset, was 94.10%, 55.18%, 93.66%, 91.33% and 68.33% respectively.

Comparing the accuracies of the best performing networks after the layer dimension permutation, with those from the result of the structure optimization (Table 1.) shows that the best accuracies grow by relatively smaller values (or no value at all), indicating that this step offers only a small improvement in the performance of the networks. The reason for the 100-sized subset of the MNIST dataset achieving the 100% accuracy is similar to as before in Table 1. However, since the final accuracy on the official MNIST 10,000-sized test set showed a rise in accuracy (the difference between full test accuracy in Table 1. and Table 2. for the MNIST, 100-sized subset), we may conclude that this step did indeed result in an increase in the performance of the network.

## Discussion

In this study, we found that primarily the sample size of the training dataset did not have a dramatic influence on the optimal network structures, and an indefinitely deeper or wider network may not necessarily be preferable. In many cases (CIFAR-10 and MNIST) performance degraded with the increase in the depth of the network. The improvement, in comparison with the average performance of a random network structure, in the classification error after optimizing the network structure was 27.99% (MNIST, 100 samples), 16.45% (MNIST, 500 samples), and 13.11% (MNIST-1000 samples). The MNIST subset with 100 samples has a much larger improvement over the average case than the 500-sized subset. Similarly, the 500-sized subset has a larger improvement over the 1000-sized subset. Comparing the improvement of the performance of the best performing structure over the average case, it can be seen that the improvement is more dramatic for subsets of smaller size and therefore our 2-step network structure search methodology is more critical in cases where small data challenges exist.

It seems that the width of each layer plays a relatively smaller role in comparison to the layer combinations, as we have seen in MNIST with 500 and 1000 samples, and CIFAR dataset with 5000 samples. However, we did observe that the influence of width could be substantial and more than other structural hyperparameters in MNIST with 100 samples and the mitosis dataset. This necessitates the optimization on both the layer configuration and the width of the layers. Furthermore, we have shown earlier that varying subsets of the same dataset can have similar optimal network structures. This potentially points to a feasible idea that network optimization may use a small subset of the entire sample to find the optimal structural hyperparameters, which can then be scaled to be trained on the entire dataset. This “in-domain” transfer learning approach uses small subsets to optimize the network structure to initialize models for a larger subset. The optimal structure can be used as a guided optimal starting point, from which training on the entire set may take place (this has been shown only on smaller sets of size < 5000 samples). This way, it is possible to find the optimal network quicker with lesser samples processed, and thus faster. This in-domain transfer learning can supplement the commonly used “cross-domain” transfer learning, in which the pre-training set and the in-training set are often of very different nature, e.g. photographic (CIFAR10) and microscopic (Mitosis) datasets. However, our results suggest that cross-domain transfer learning will be most useful when the nature of the data in the pre-training set, and the ad-hoc set is similar. This means that it may not always work if the data nature is very different because the optimal networks are intensely data-driven and each set may have very contrasting optimal structures. In this case, using networks which have been pre-trained (and hence show optimal performance on a particular set) may not necessarily reflect the same optimal performance in the ad-hoc set. Nonetheless, transfer learning itself particularly when and in what application exactly pre-training works or how pre-training should be applied (in-domain, cross-domain, fine-tuning or feature extraction) is still something that needs to be further studied.

In the field of medical imaging specifically, lack of sufficient data and skewed datasets (like the Mitosis dataset) can lead to underperforming networks in the case when those networks are organized using a generalized rule, with the hope of achieving the highest performing network. In this case, using a subset of data for structural optimization would offer a more guided approach to acquiring the most optimal network. This may help with taking into consideration the lack of data/skewed nature of the data (by recording their lower performance) and provide an efficient arrangement and number for the layers as well as an optimal width. There are several publicly available imaging archives that have less than 1,000 samples. The Grand Challenge for Breast Cancer Histology images (https://iciar2018-challenge.grand-challenge.org/home/) involving automatically classifying H&E stained breast histology microscopy images into four classes: normal, benign, in-situ carcinoma and invasive carcinoma, contain only 400 microscopy images, with only 100 samples per class. The Diabetic Retinopathy Segmentation and Grading Challenge (https://idrid.grand-challenge.org/home/) involving tasks such as lesion segmentation, disease grading and optic discs and fovea detection contain only around 516 images in the dataset, with an extremely high dimensionality of 4288×2848 pixels per image. We have demonstrated our approach using the Mitosis dataset in this study, and it can also be applied to the above-mentioned data sets, which have similar small data challenges.

There are limitations to this study. Due to the use of a tree structure to enumerate the list of network structures within a constrained space, the number of such structures grows exponentially with the increase in the maximum allowed VC-dimension of the structures. This can result in an arbitrarily large list of structures which may take very long to train and validate. It is possible that we can miss a good performing network that has large VC-dimension. The same issue lies with the layer dimension permutation enlisting for a particular structure. The number of possible layer dimension permutations grows exponentially with the depth of the structure. The limitations in both cases are due to the number of permutations blowing up as the number of paths from the root to the leaves of the tree structure grows exponentially with the height (in this case) and width of the tree. A possible follow-up solution for this limitation would be to have some form of tree-pruning or permutation/structure dropping algorithm to remove candidate structures before training and validation which are known to either be too identical in performance to other structures or are known beforehand to be sub-optimal structures.

Another limitation in this study is that the cross-validation techniques were simple dataset splits followed by per epoch validation (Holdout method) and did not use any sophisticated validation techniques like K-fold [17], Stratified K-fold, Leave-P-Out, etc. In certain cases, where datasets are much smaller (like MNIST-100), the use of more complex validation methods may be more suitable. Since we needed to keep all the training/validation parameters and methods consistent across all sets we maintained the trivial cross-validation method for all datasets/subsets. Through the similar idea, various other optimizers and objective functions may be used, depending on the suitability of the problem/dataset.

One possible improvement in this study is to use a more rigorous approach to select the subset of the entire dataset for structural optimization. When using a smaller subset of a larger dataset, like in the case of MNIST and CIFAR-10, to find the optimal network, the selection process for the subset samples was done randomly. Consequently, we may require larger subsets to adequately train the networks to take into consideration samples which are highly correlated. In this case a sample pre-selection algorithm could be designed, to test the correlation between two samples and therefore select only one, For example, we may do sampling over clustered data samples to de-correlate the sampled subset. This would help to remove redundancy in the subset and consequently allow the subset to be smaller, hence allowing the training and validation to complete quicker. Additionally, in many cases, because the number of networks maybe large (order of 1000), the training and validation of the two steps (structure selection and layer dimension permutation) may take a long time to run. Again, this can be sped up using high-performance clusters or certain optimizations as deemed suitable for the ad-hoc practice, like using distributed computing, as each structure’s training and testing is independent and suitable for parallelism.

In conclusion, our study shows that the optimally performing network is largely determined by the data nature, and the data size plays a relatively much smaller role. For a sufficiently small sample size, a separate network structure optimization step, along with a layer dimension optimization step can be a useful strategy to find the optimally performing network as the two-step heuristic offers a more exhaustive approach to the optimization of the best performing network.

## References

[1] A. Krizhevsky, I. Sutskever, and G. E. Hinton, “Imagenet classification with deep convolutional neural networks,” in Advances in neural information processing systems, 2012, pp. 1097–1105.

[2] K. Simonyan and A. Zisserman, “Very deep convolutional networks for large-scale image recognition,” arXiv preprint arXiv:1409.1556, 2014.

[3] C. Szegedy et al., “Going deeper with convolutions,” in Proceedings of the IEEE conference on computer vision and pattern recognition, 2015, pp. 1–9.

[4] K. He, X. Zhang, S. Ren, and J. Sun, “Deep residual learning for image recognition,” in Proceedings of the IEEE conference on computer vision and pattern recognition, 2016, pp. 770–778.

[5] M. Oquab, L. Bottou, I. Laptev, and J. Sivic, “Learning and transferring mid-level image representations using convolutional neural networks,” in Computer Vision and Pattern Recognition (CVPR), 2014 IEEE Conference on, 2014, pp. 1717-1724: IEEE.

[6] J. Donahue et al., “Decaf: A deep convolutional activation feature for generic visual recognition,” in International conference on machine learning, 2014, pp. 647–655.

[7] B. Q. Huynh, H. Li, and M. L. Giger, “Digital mammographic tumor classification using transfer learning from deep convolutional neural networks,” Journal of Medical Imaging, vol. 3, no. 3, p. 034501, 2016.

[8] N. Tajbakhsh et al., “Convolutional neural networks for medical image analysis: Full training or fine tuning?,” IEEE transactions on medical imaging, vol. 35, no. 5, pp. 1299–1312, 2016.

[9] Y. LeCun, C. Cortes, and C. Burges, “MNIST handwritten digit database,” AT&T Labs [Online]. Available: http://yann.lecun.com/exdb/mnist, vol. 2, 2010.

[10] A. Krizhevsky, V. Nair, and G. Hinton, “The CIFAR-10 dataset,” online: http://www.cs.toronto.edu/kriz/cifar.html, 2014.

[11] V. Vapnik, E. Levin, and Y. L. Cun, “Measuring the VC-dimension of a learning machine,” Neural computation, vol. 6, no. 5, pp. 851–876, 1994.

[12] E. D. Sontag, “VC dimension of neural networks,” NATO ASI Series F Computer and Systems Sciences, vol. 168, pp. 69–96, 1998.

[13] N. Harvey, C. Liaw, and A. Mehrabian, “Nearly-tight VC-dimension bounds for piecewise linear neural networks,” arXiv preprint arXiv:1703.02930, 2017.

[14] D. Kingma and J. Ba, “Adam: A method for stochastic optimization,” arXiv preprint arXiv:1412.6980, 2014.

[15] L. Prechelt, “Automatic early stopping using cross validation: quantifying the criteria,” Neural Networks, vol. 11, no. 4, pp. 761–767, 1998.

[16] F.-C. Yeh, Q. Ye, T. Hitchens, Y. Wu, A. Parwani, and C. Ho, “Mapping stain distribution in pathology slides using whole slide imaging,” Journal of Pathology Informatics, Research Article vol. 5, no. 1, pp. 1–1, January 1, 2014 2014.

[17] T. Fushiki, “Estimation of prediction error by using K-fold cross-validation,” Statistics and Computing, journal article vol. 21, no. 2, pp. 137-146, April 01 2011.

